# Integrating Transmission Dynamics and Pathogen Evolution Through a Bayesian Approach

**DOI:** 10.1101/2024.04.15.589468

**Authors:** Ugnė Stolz, Tanja Stadler, Timothy G. Vaughan

**Affiliations:** Department of Biosystems Science and Engineering, ETH Zürich, Mattenstrasse 4058 Basel, Switzerland; Swiss Institute of Bioinformatics (SIB), 1015 Lausanne, Switzerland

## Abstract

The collection of pathogen samples and subsequent genetic sequencing enables the reconstruction of phylogenies, shedding light on transmission dynamics. However, many existing phylogenetic methods fall short by neglecting within-host diversity and the impact of transmission bottlenecks, leading to inaccuracies in understanding epidemic spread. This paper introduces the *Transmission Tree (TnT)* model, which leverages multiple pathogen gene trees to more accurately model transmission history. By extending the Bayesian phylogenetic analysis software *BEAST2, TnT* integrates the sampled ancestor birth-death model for transmission trees and the multi-species coalescent model for pathogen gene trees. This integration allows for the consideration of critical factors like transmission orientation, incomplete lineage sorting, and within- and between-host diversity. Notably, *TnT* incorporates an analytical approach to address unobserved transmission events, crucial in scenarios with incomplete sampling. Through theoretical evaluation and application to real-world cases like HIV transmission chains, we demonstrate that *TnT* offers a robust solution to improve understanding of epidemic dynamics by effectively combining pathogen gene sequences and clinical data.

## 1 Introduction

Disease surveillance involves the collection and sequencing of pathogen samples that enable the reconstruction of phylogenies. Such phylogenies often contain useful information about the transmission dynamics that gave rise to the observed sequence diversity. A variety of phylogenetic methods for learning about transmission dynamics exist and it is common for most of them to assume perfect identity between pathogen phylogenies and transmission trees, ignoring potential disparities between within- and between-host transmission dynamics (Romero-Severson et al. 2014). In particular, this means that most of them use a single quasispecies or consensus sequence per host. Particularly in the presence of high within-host variability and strong transmission bottlenecks, this identity assumption disregards key factors like incomplete lineage sorting, transmission direction, within-host diversity, the strength of transmission bottlenecks (innoculum size), and the diverse nature of host sampling schemes. These factors are, however, critical when assessing transmission trees, since incomplete lineage sorting produces differences between the pathogen phylogenies and transmission trees with respect to the order and relatedness of the transmission pairs. Determining the correct transmission direction and strength of the bottleneck is essential to accurately characterize within-host quasispecies and discover the seeding viral variant (Zwart and Elena 2015; Joseph et al. 2015). Furthermore, simplistic host sampling schemes that allow complete sampling or impose a single sampling timepoint per host are unrealistic for most transmission clusters and may discard available data.

There have been several recent attempts to address the above issues related to the simplistic identity assumption and disregard of key factors, offering varying degrees of improvement. We briefly review some of the most well-known models outlining their key features, as well as disadvantages, we aim to overcome in this paper. The *outbreaker2* (Campbell et al. 2018) is based on genetic distances and utilises contact tracing information but without fairly complicated customization it does not account for within-host diversity. While the program is modular and a custom model can be implemented, it requires some programming knowledge, and thus might not be accessible to every user. *SCOTTI* (Maio, Wu, and Wilson 2016), employing a structured coalescent approach to model transmission as viral migration between hosts, allows incomplete host sampling and multiple samples per host. However, it requires user-defined exposure intervals during which a host can be part of an epidemic, and does not allow for transmission bottlenecks. *TransPhylo* (Didelot et al. 2017; Didelot et al. 2021; Carson et al. 2024) can handle incomplete sampling of infected individuals but requires the input of a pre-estimated phylogenetic tree of the pathogen. The *phybreak* (Klinkenberg et al. 2017) package circumvents this by attempting simultaneous transmission and viral gene tree inference and accounting for within-host variation and transmission bottleneck. Still, it does not explicitly model unobserved transmission events and thus is only suitable for densely sampled outbreaks. To ensure applicability in a variety of scenarios and to avoid constraints that stem from more complicated priors, *phybreak* assumes simplistic transmission tree generation and sampling models, which exclude some epidemiologically meaningful parameters. The *BadTrIP* (Maio et al. 2018) package makes use of phylogenetic models with polymorphisms (Maio, Schrempf, and Kosiol 2015) and can directly employ next generation sequencing (NGS) data. It accounts for transmission bottlenecks, but in a rigid manner where a discrete number of lineages passing though upon a transmission event is determined. *BadTrIP* also assumes complete sampling and cannot efficiently handle datasets exceeding 100 samples. A faster method *TNet* (Dhar et al. 2020), which applies Sankoff’s algorithm to maximum likelihood phylogenies, can handle thousands of NGS sequences and accounts for within-host diversity. It does not explicitly model the bottleneck and was tested only on simulations where full sampling is assumed. Finally, there have been recent advancements in highly scalable models for transmission network inference, when taking into account the within host pathogen diversity (Skums et al. 2022; Specht et al. 2023; Carson et al. 2024). However, they either require a time scaled phylogeny as an input (Skums et al. 2022; Carson et al. 2024), thus potentially limiting the full exploration of phylogenetic uncertainties, or are non-tree based (Specht et al. 2023). Moreover, a common limitation among these models is their inability to explicitly model unobserved transmission events or to accurately infer the timing of such events. This gap underscores a need in transmission inference: a model that can account for transmission bottleneck, complete transmission history, and within-host diversity, as well as provide uncertainty intervals for its estimates. See also recent work by Duault, Durand, and Canini 2022 and Sobkowiak et al. 2022 for a more detailed review and comparison of the available methods.

In this paper, we propose a new approach that uses a framework which estimates multiple congruent trees—- both transmission and pathogen gene trees—to model the evolutionary history more accurately. This method builds upon and extends the principles used in macroevolutionary studies for joint species and gene tree inference (Heled and Drummond 2009; Ogilvie, Bouckaert, and Drummond 2017; Douglas, Jiménez-Silva, and Bouckaert 2022). Specifically, we employ the sampled ancestor birth-death (SABD, Stadler 2010; Heath, Huelsenbeck, and Stadler 2014; Gavryushkina et al. 2014) model to describe the transmission tree and extend the multi-species coalescent (MSC, Degnan and Rosenberg 2009) to include non-deterministic transmission bottlenecks for the corresponding viral gene tree inference. This combination allows us to account for transmission orientation, incomplete lineage sorting, and the within- and between-host diversity. Additionally, we employ an analytical approach to integrate over all unobserved transmission events, facilitating transmission inference in situations with incomplete sampling. We detail the development and validation of our package, named *TnT*, which extends the Bayesian phylogenetic analysis software *BEAST2*. We demonstrate its utility and advantages over existing models through a rigorous theoretical and practical evaluation, including a comparison with the *SCOTTI* model on simulated data and the application of *TnT* to a known HIV transmission chain.

## 2 Materials and Methods

### 2.1 Phylodynamic Model

We use and adapt the Bayesian framework presented by Heled and Drummond (2009), Ogilvie, Bouckaert, and Drummond (2017), and Douglas, Jiménez-Silva, and Bouckaert (2022), which was developed for the macroevolution context. Specifically, we employ a Markov chain Monte Carlo (MCMC) algorithm to effectively integrate over the pathogen gene trees and obtain the marginal posterior distribution over the host transmission tree *T*, given sequence alignments for each of *n* loci *A* = *a*_1_, *a*_2_, …, *a*_*d*_, as described by equation (1) in Heled and Drummond (2009):

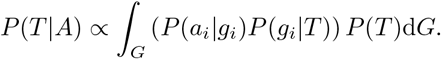

Here *G* = (*G*_1_ × *G*_2_ × … × *G*_*d*_) is the space of all gene trees and *g*_*i*_ ∈*G*_*i*_ a specific gene tree for *i*^*th*^ alignment.

One key difference from the previous work is the inclusion of transmission direction in our model, informed by sampling information, such as host-sample allocation and sampling dates. Another is our inclusion of a transmission bottleneck, which is modelled as an instantaneous event at transmission. Every bifurcation on the transmission tree is an observed transmission event, while unobserved events are allowed along the branches. The latter ensures the possibility of incomplete sampling. This tree is best visualised as an asymmetric tree where the lineage that started before the bifurcation event and continues afterwards represents a *donor* host and the lineage that branches from it at said event is a *recipient* host. Figure 1 illustrates the congruent transmission and gene trees in the *TnT* model, along with its main features.

**Figure 1.**
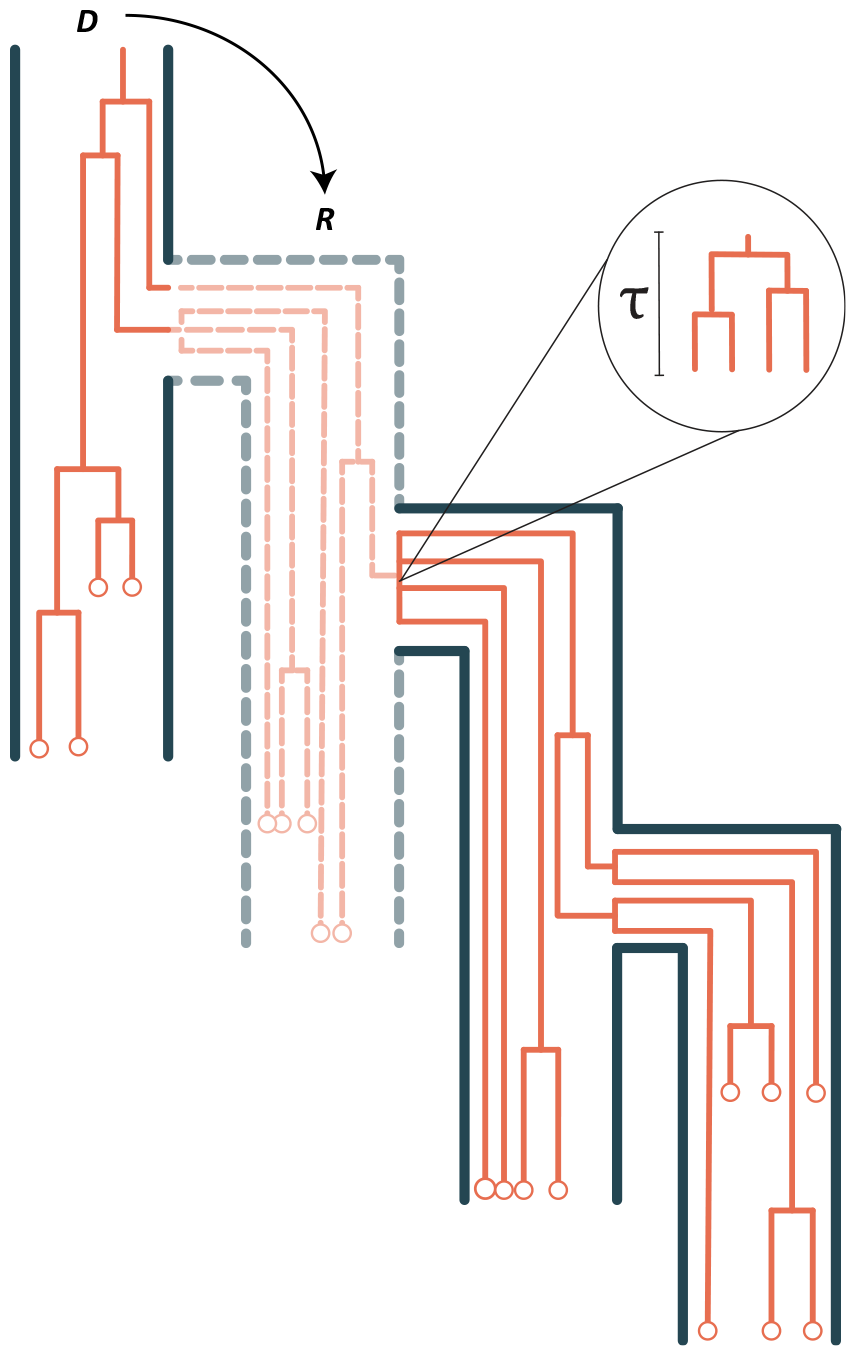
Figure shows the embedding of a gene tree (orange) within the oriented transmission tree (dark contour) and all possible events. For simplicity, a single gene tree is shown, however, there may be many. The empty circles show gene samples, which can be collected through time. We orientate this tree by assigning donor (**D**) and recipient (**R**) label at each observed transmission event. We leave the donor on the left-hand side and recipient on the right. The faded dashed branch shows an unsampled and hence unobserved host in the transmission chain. At any (observed or unobserved) transmission event, bottleneck may induce multiple instantaneous coalescent events that show up as bifurcating, multifurcating, or multiple merger events. Under some effective population size, we may expand this instantaneous event and obtain the duration *τ*, needed for it to happen.

#### 2.1.1 Transmission Bottleneck as an Instantaneous Event

We model the instantaneous transmission bottleneck that occurs at the end of the recipient edge (looking from present to past) during every transmission event, irrespective of whether it is observed or unobserved. The bottleneck is parameterised by the probability *p* := 1 − *e*^*−τ*^, which represents the probability that two gene tree lines will coalesce at the same transmission event. Here, *τ* is the time required to observe the equivalent amount of coalescence under the prevailing effective population size (Figure 1, see the Supplementary Material for more details on the bottleneck strength definition). Bottlenecks can induce coalescent, multifurcation, or multiple merger events on the gene tree.

A *bifurcation* event corresponds to the merging of exactly two lineages at their common ancestor. A *multifurcation* event involves the joining of more than two lineages at a single common ancestor. A *multiple merger* is an event in which two or more bifurcations or multifurcations occur simultaneously. We will refer to the latter two possibilities as *higher-order coalescence*. In instances of a weak bottleneck or when only one lineage, ancestral to our samples, passes through the bottleneck, we may not observe any events at transmission.

The instantaneous bottleneck idea as part of the coalescent process has previously been described in detail by Galtier, Depaulis, and Barton 2000. It has also been recently implemented by Baumdicker et al. 2022 for coalescent simulations.

#### 2.1.2 Unobserved Transmission Events Follow a Poisson Distribution

Take a birth-death-sampling process with probabilistic removal on sampling, where we denote birth rate *λ*, death rate *µ*, sampling through time rate *ψ*, removal probability *r* and sampling at present probability *ρ* and ensure all rates to be positive and *λ* ≠ *µ*. Following Stadler (2010) and Gavryushkina et al. (2014), the probability that an individual alive at time t before today has no sampled extinct or extant descendants is:

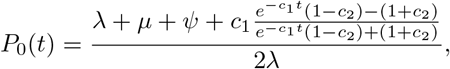

where

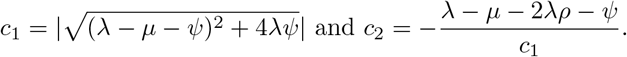

**Table 1:**
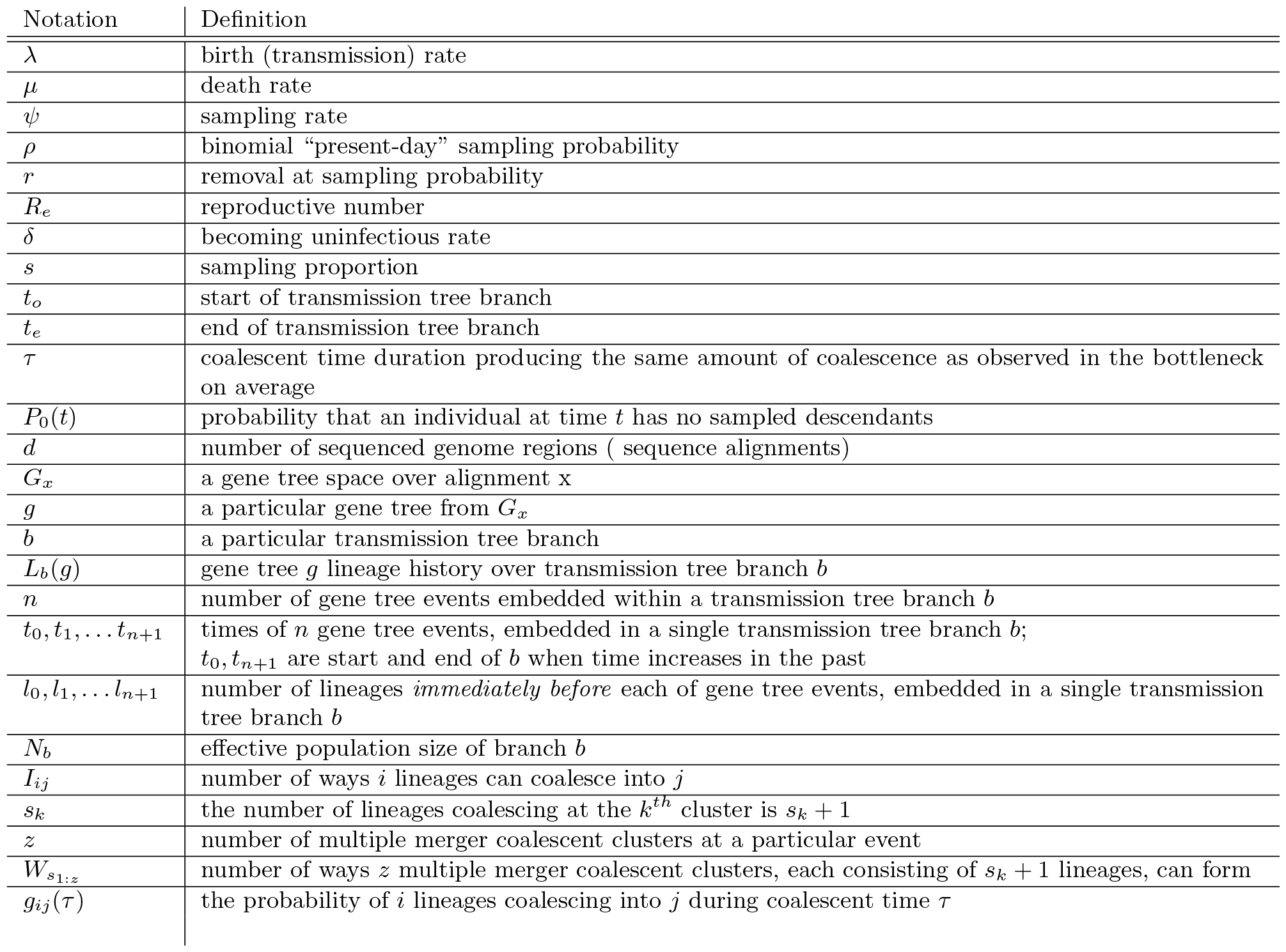
Notation used in this paper.

Define also

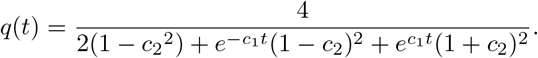

Following Bokma, Brink, and Stadler (2012), the chance of unobserved bifurcation is 2*λP*_0_. Then the expected number of unobserved bifurcations on a branch between its origin time *t*_*o*_ and end time *t*_*e*_, when time increases backwards, is

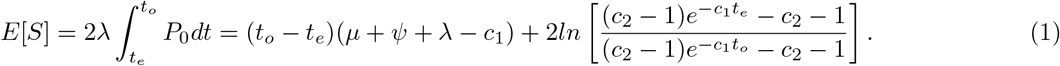

Further, we are interested in those unobserved transmission events that *occur along a branch*. That is, immediately before an unobserved transmission event on a branch, the host associated with this branch is the donor, and immediately after this event it is the recipient. We call such events hidden transmission events on the tree. It can be shown that the number of such events between time *t*_*o*_ and *t*_*e*_ follows a Poisson distribution with mean 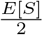 (see the Supplementary Material for derivation). The times of hidden transmission events that change the host of a branch follow a non-homogeneous Poisson process with rate *λP*_0_.

#### 2.1.3 Jointly Modelling the Transmission and Pathogen Evolution

##### Transmission Tree Prior

We model the transmission tree as a oriented birth-death process and allow for sampled ancestors, present day sampling and sampling through time (Stadler 2010; Stadler et al. 2012; Gavryushkina et al. 2014). We rely on the implementation of the sampled ancestor birth death (SABD) model, found in the BDMM-Prime (Vaughan 2022) package, as input to *TnT*.

For easier interpretation, the SABD model can be parameterised in different ways (Gavryushkina et al. 2014).

In our simulations and data analysis, we will use the epidemiological parameterisation with effective reproductive number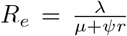, becoming uninfectious rate *δ* = *µ* + *ψr*, sampling through time proportion 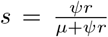 and removal at sampling rate *r*.

##### Gene Tree Prior

Assume that we have *d* distinct pathogen genome regions sequenced. These may represent genes or segments (in segmented viruses) or differently defined regions of interest in the pathogen genome. We then model *d* trees, which are informed by these alignments, by the coalescent process with instantaneous bottlenecks. We call them *gene trees* and restrict each of them by the same transmission history as modelled by the transmission tree.

Let *g* be a gene tree from the space of gene trees *G*_*x*_ over the *x*^*th*^ alignment (1 ≤*x* ≤ *d*). Moving backward in time along the branch *b* of the transmission tree, we can observe all events on this embedded gene tree and their respective times *t*_0_, *t*_1_, …, *t*_*n*_, *t*_*n*+1_. Here, times *t*_0_ and *t*_*n*+1_ represent the start and end of the branch *b*. Similar to Heled and Drummond (2009), we denote the gene tree lineage history over branch *b* as *L*_*b*_(*g*) = {*l*_*m*_, *t*_*m*_|0 ≤ *m* ≤ *n* + 1}. Here, *l*_*m*_ is the number of tree lineages at time *t*_*m*_. Unlike Heled and Drummond (2009), the number of lineages at time *t*_*n*+1_ depends not only on coalescent and sampling events. It can also be affected by bifurcations, multifurcations or multiple mergers, introduced by instantaneous bottlenecks at the hidden transmission events along the branch *b*.

Furthermore, if the branch *b* of the transmission tree corresponds to a donor host, there are no events on a gene tree *g* at the time *t*_*n*+1_. Otherwise, if it corresponds to a recipient host, we model an instantaneous transmission bottleneck at time *t*_*n*+1_. It is worth noting that the bottleneck increases the probability of two lineages to coalesce, going backwards in time, but it does not imply that bifurcation, multifurcation, or multiple merger event will necessarily be observed on any gene tree.

We define the probability of a gene tree embedded within a transmission tree branch *b*, having a history *L*_*b*_ as the product of contributions from bifurcating and higher-order coalescent events, as well as the intervals between them. Each bifurcation, in turn, can account for a coalescent event, induced by the usual coalescent process, or a coalescent event resulting from a transmission bottleneck at a transmission event (hidden or observed). Each higher-order coalescence, reducing *i* lineages to *j*, can only be the result of an instantaneous bottleneck and therefore must be a result of a transmission event. It is important that a hidden transmission event may also occur without inducing any events on a gene tree, either because there is only one gene tree lineage that is involved in this event and ancestral to the samples at the tree tips, or because the bottleneck is too weak.

In summary, the multispecies coalescent likelihood of a gene tree can be decomposed as a product of likelihoods over the transmission tree branches *b* (Heled and Drummond 2009). In order to account for the bottleneck at observed and unobserved transmission events, we have to modify equation (2) of Heled and Drummond (2009). That is, going through all *n* + 1 intervals and events for a transmission tree branch we have:

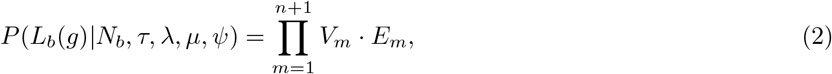

where *V*_*m*_ is the *m*^*th*^ interval contribution and *E*_*m*_ is the corresponding event ending the interval. We define the interval contributions as:

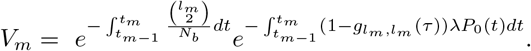

In order to write down the event contribution, we will use a helper function *h*(*τ, i, j, z*), where *i* is the number of lineages before the event, *j* is the number of lineages after the event, *z* is the number of coalescent clusters that make up the event and the set **S** = { *s*_*k*_ | *s*_*k*_ ∈ ℕ ∧*s*_*k*_ *>* 0 ∧1 ≤*k* ≤*z*}, where *s*_*k*_ + 1 is the number of lineages that coalesce in a cluster *k*:

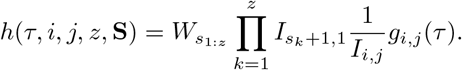

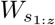 denotes the number of ways *z* multiple merger groups, each consisting of *s*_*k*_ + 1 lineages, can form, *I*_*ij*_ denotes the number of ways *i* lineages can coalesce into *j* and *g*_*ij*_(*τ*) is the probability of *i* lineages coalescing into *j* during coalescent time *τ*.

In case of bifurcation, *z* = 1 and the number of lineages after the event are one less than before: *j* = *i* − 1. Also,

**S** = {*s*_1_ = 1}, giving, 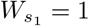 and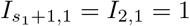. Then, we have

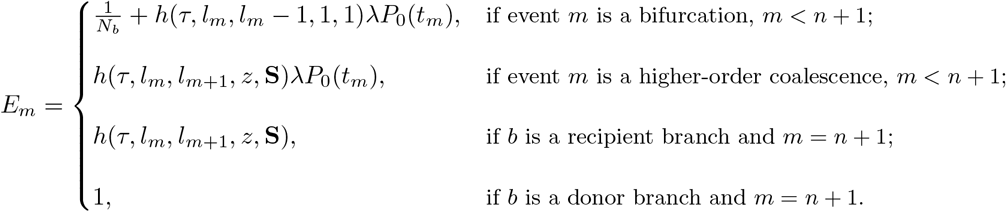

Here, *N*_*b*_ is the effective population size of branch *b* and *λP*_0_(*t*) is the rate at which unobserved transmission events that switch the host occupying a branch happen. See the Supplementary Material for the derivation.

### 2.2 Stochastic Mapping of Unobserved Transmission Events

Given a posterior sample, consisting of a transmission tree and its associated parameters of the SABD model, we are able to use the stochastic mapping technique (Nielsen 2002; Huelsenbeck, Nielsen, and Bollback 2003) to recover the unobserved transmission events that changed host occupation along the tree branch. Since these events follow a Poisson process with a known rate, we may apply the inverse transform sampling. Further, we only need to sample the timing of the events and not the host type as we do not care which unobserved host occupied the branch, only that the transmission happened.

As long as the posterior trees and parameter estimates are recorded, stochastic mapping may be performed after the analysis is complete. Since in the current version we are ignoring the events on the gene trees, it provides us with full but approximate transmission history between the observed samples. Therefore, even in the lack of full sampling we are able to quantify the transmission times for each observed host.

### 2.3 Validation

To validate *TnT* and ensure that the parameters are well recovered, we performed a well-calibrated study (Dawid 1982). We have simulated 200 SABD transmission trees with a single embedded gene tree. Host samples were collected heterochronically and each contained 5 contemporaneous within-host pathogen samples. We also conditioned the transmission trees to be between 5 and 30 observed hosts (with total number of hosts unrestricted) for a more manageable analysis. Parameters were independently drawn from the following distributions: *R*_*e*_ ∼Uniform(1, 4), *δ* ∼Uniform(0.1, 2.0), *s* ∼Uniform(0.3, 1.0), *r* ∼Uniform(0.2, 1.0), *Ne* ∼LogNormal(1, 0.5), *p* ∼Uniform(0, 1), *ρ* ∼Uniform(0, 1). Transmission process origin was always assumed to be 3 time units in the past. Finally, we simulate genetic sequences along these gene trees using the strict clock with rate 0.005 and the Jukes-Cantor substitution model.

Then, we conducted inference for each transmission and gene tree pair using these sequences and sample times as data. We used the above true distributions as parameter priors for all parameters, except for sampling proportion, which was assumed known. We assumed that the true clock and substitution models were known.

Data simulation and analysis were performed by the following R R Core Team 2022 packages: beastio (Plessis), ape (Paradis, Claude, and Strimmer 2004), phytools (Revell 2012), ggplot2 (Wickham 2016), coda (Plummer et al. 2006) and stringr (Wickham 2023) (see Data and Software Availability).

### 2.4 Assessing Inference Power of *TnT* and *SCOTTI* on Simulated Data

While the binomial “present-day” sampling probability *ρ* can be estimated as part of the model, this is less useful in epidemiological studies, since it is rarely assumed that pathogen sequences are collected at a single time. Thus, we fix this parameter to zero in what follows. Further, in order to evaluate not only the recovery of model parameters but also transmission traits such as host infection time and number of unobserved intermediate hosts, we need to know the full transmission history. Therefore, for further analysis, we simulated additional 100 full BD transmission histories with a single embedded gene tree. Then, we subsampled these histories over time to obtain the observed dataset. We have used the same distributions as above to draw parameters, except for *ρ* = 0. We again had 5 pathogen samples per host sample, but allowed 5 to 50 hosts samples in each SABD transmission tree. Genetic sequences along the gene tree were simulated as above.

We used these observed samples as data for *TnT* and *SCOTTI* and full transmission histories to evaluate their performance. We performed *TnT* analyses as described above in the Validation section. Here, we do not model sampling at present, therefore we also provided true sample offset value, which notes how far the last sample is from the present. For *SCOTTI*, which uses structured coalescent as a transmission framework, we retained the default uninformative effective population size and migration rate priors. We again used the true clock and substitution models. Since the simulation model does not explicitly model exposure intervals, for *SCOTTI* analyses, we assumed uninformative exposure times for each host. We bounded the total number of hosts (observed and unobserved) in the chain to 1000 (which is more than the maximum number obtained in any of the simulations).

#### Application to a Well-Documented HIV Chain

We applied *TnT* to a HIV-1 subtype C patient transmission chain involving 11 hosts that was previously extensively described (Lemey et al. 2005; Vrancken et al. 2014). We used the 449 available *env* (212) and *pol* (237) sequences with drug resistance associated targets removed from *pol* alignment as in Vrancken et al. (2014). The prior parameter distributions were *R*_*e*_ ∼LogNormal(0.5, 1.0), *δ* ∼LogNormal(− 1.0, 1.0), *s* = 1.0, *ρ* = 1.0, *r* ∼Beta(5.0, 2.0), *Ne* ∼LogNormal(0.44, 0.5), *p* ∼Beta(3, 1.5), origin ∼LogNormal(3, 0.1). Due to the extensive clinical data, we assume complete sampling for this chain, which is ensured by values of sampling through time proportion *s* and sampling at present probability *ρ*. By “present”, here we mean the time of the last sample collection. This is necessary to ensure that no unobserved hosts are modeled within this chain. We did not assume a different effective population size above the origin. Lastly, we estimate the rates of the strict clock and substitution model separately for both genes and use the general time-reversible substitution model with four gamma rate categories (GTR + Γ). We ran 3 identical MCMC chains with different random starting values and analysed the combined chain after removing 30% burn-in. In addition, we repeat the inference with exactly the same set up, except that no bottleneck is assumed. These chains converged faster and we combined them after removing 20% burn-in. See Supplementary Figure 1 for the trace and effective sample size (ESS) of both analyses.

## 3 Results

### 3.1 Inference of Transmission Parameters and Times

The well-calibrated study is summarised in Figure 2, which shows that the parameters for both transmission and gene trees were accurately retrieved. As expected the fraction of times a given parameter true value falls within highest posterior density (HPD) credible interval is approximately equal to the relative width of this interval.

**Figure 2.**
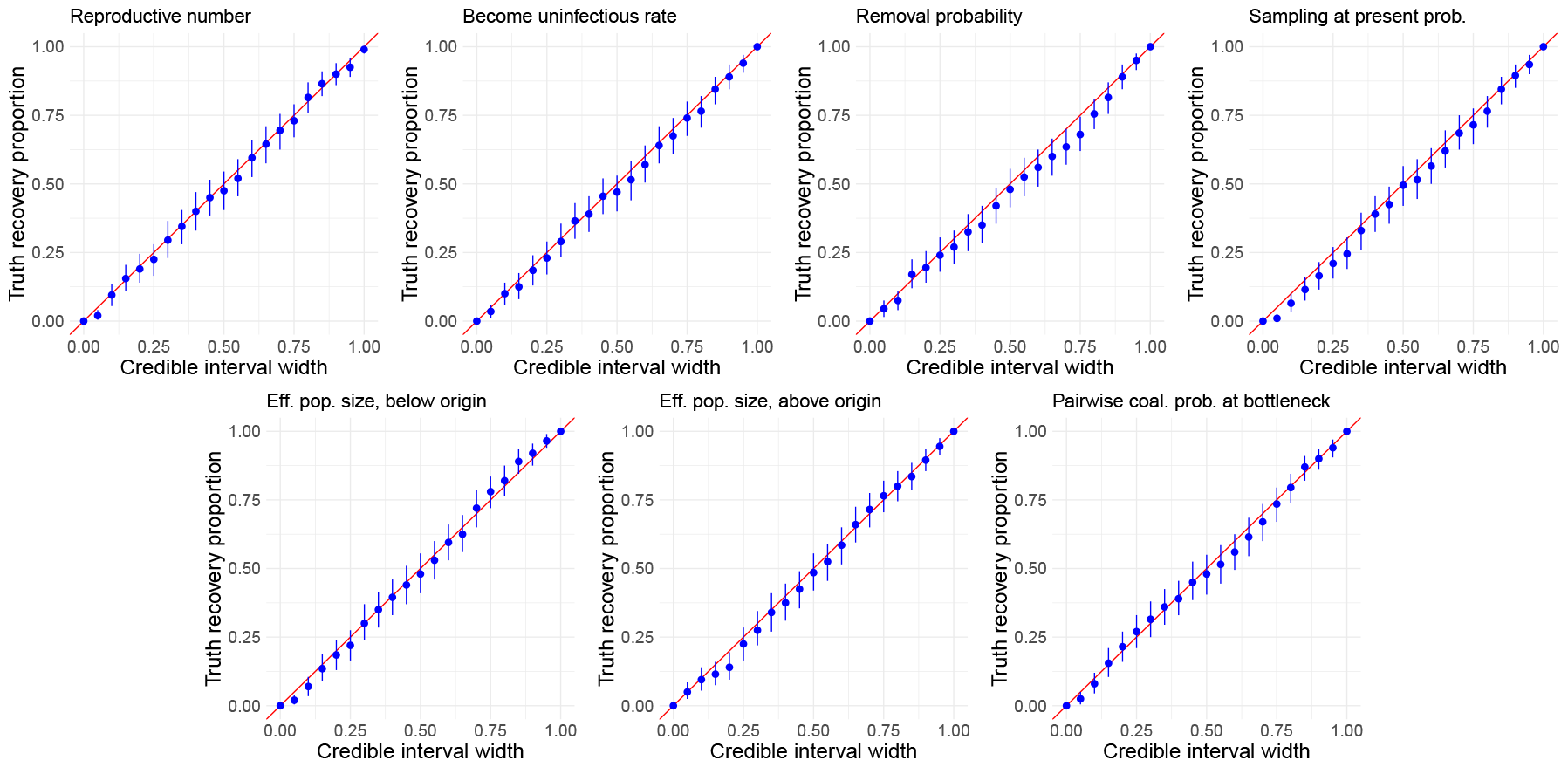
Results of the well calibrated simulation study. The dots correspond to the proportion of simulations where the true value fell within the HPD credible interval of a certain width. The vertical lines show bootstrap confidence intervals. The diagonal x = y shows the expected result.

When performing the second set of simulations, we recorded not only the sampled transmission trees, but also the complete transmission history including transmission times. This allows us to know when exactly a host in the sampled tree was infected, even when the direct transmitter was not sampled. In order to recover the full history on the inference side, we performed stochastic mapping on the posterior transmission trees. Then we compared the true and inferred infection times and showed that they are well recovered: the fraction of true values within the 95% HPD interval was 0.96 for unobserved and 0.87 for observed transmissions (Figure 3a). Unobserved transmission times had larger HPD intervals with an average relative HPD width of 2.8, compared to 0.9 for observed transmissions.

**Figure 3.**
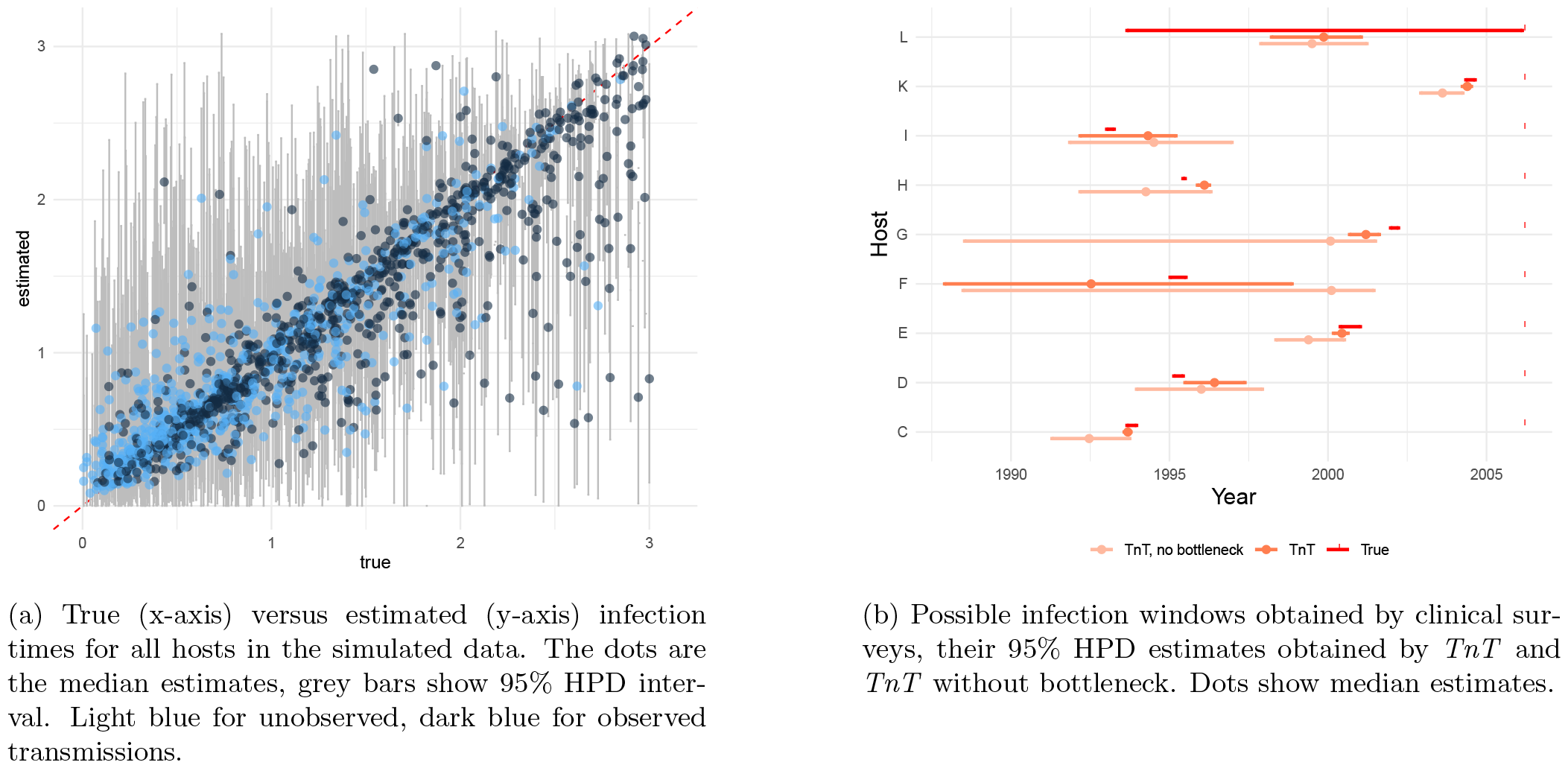
Inference of infection times, obtained by *TnT*, on simulated and real data.

### 3.2 TnT Outperformed SCOTTI on Simulated Data

Next, we investigate the performance of *TnT* compared to *SCOTTI*. While *SCOTTI* is also a Bayesian mode, its assumptions are very different. It is based on a single-tree coalescent analysis and treats transmission events as migrations between different hosts. It allows for unsampled hosts and double infections but does not account for a transmission bottleneck or estimate infection times.

Since it is not clear how best to compare the models on indirect observable transmission events (where some intermediate hosts are not sampled), we compare only direct transmission events (when both transmitter and recipient are sampled) for all tree posterior sets in the simulated data. For each simulation, both methods can quantify the posterior probability of directed transmission for any pair of hosts. Therefore, we can view this as a classification problem and apply the usual comparison metrics for such problems. Figure 4 shows the receiver operator curve (ROC) and the precision recall curve (PRC). We can see in the ROC that while both models achieve a high area under the curve, *TnT* clearly outperforms *SCOTTI*. However, since the data is very sparse (many pairs of hosts will have a direct transmission probability of 0), the PRC is a better performance metric (Davis and Goadrich 2006). This is well-illustrated by the grey dashed line in Figure 4, which depicts the performance of a random classifier.

**Figure 4.**
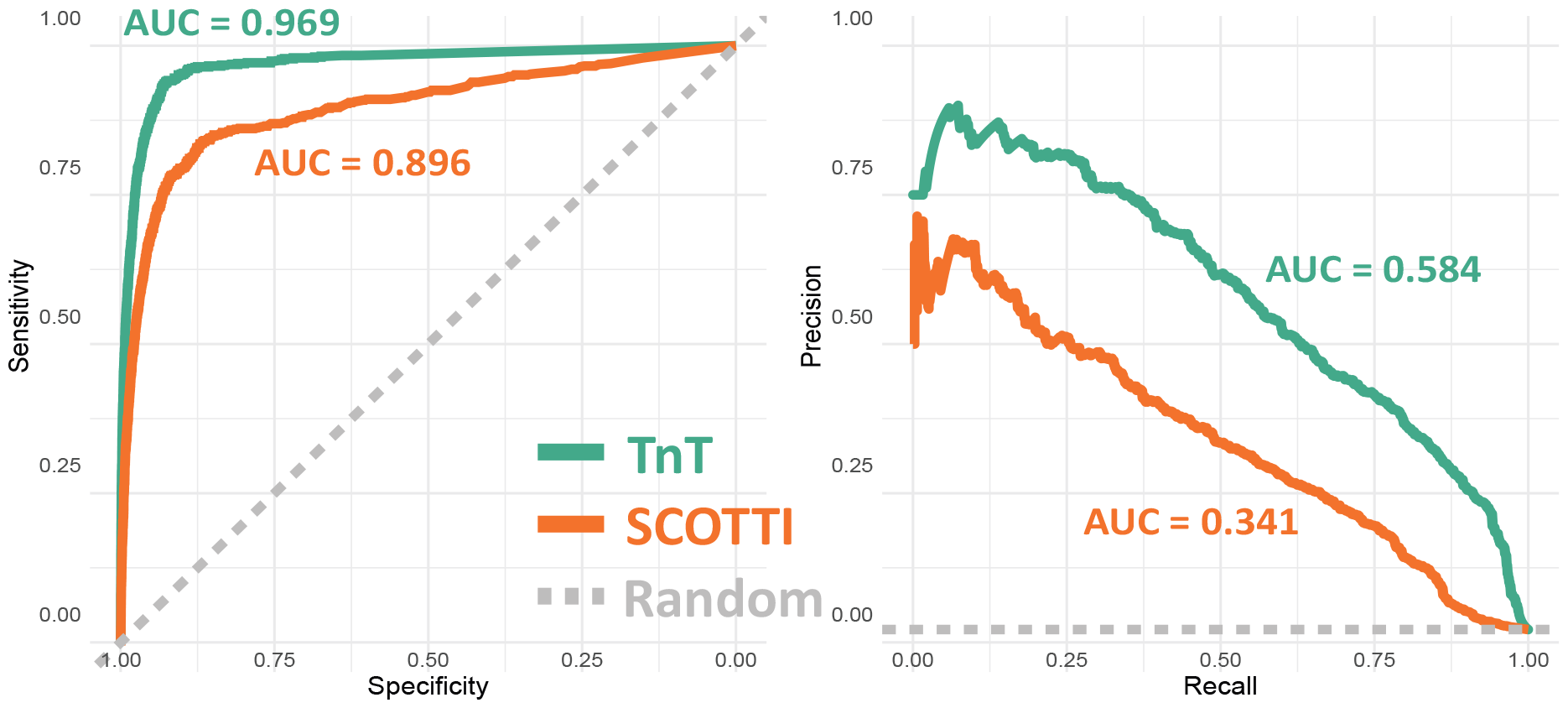
Comparison of the *direct* transmission events in the simulated dataset between *TnT* (green) and *SCOTTI* (orange) and a random classifier. **Left** ROC curve shows a better performance of *TnT*. **Right** The precision-recall curve shows better performance of *TnT*.

While these results usefully evaluate the overall performance of both models in identifying transmission pairs, practical application of either method requires definition of a probability threshold at which direct transmission events are to be considered “real”. Table 2 shows the classification results for *TnT* and *SCOTTI* with a threshold of 0.5. In particular, *TnT* achieves higher sensitivity and positive predictive value than *SCOTTI*, while specificity and negative predictive value are comparable.

**Table 2:** Performance of *TnT* and *SCOTTI* on direct transmission events in simulated data, when we apply probability threshold of 0.5 in transmission pairs classification. PPV – positive predictive value; NPV – negative predictive value.

|  | TnT | SCOTTI |
| --- | --- | --- |
| Sensitivity | 0.454 | 0.373 |
| Specificity | 0.995 | 0.989 |
| PPV | 0.663 | 0.421 |
| NPV | 0.988 | 0.986 |

### 3.3 TnT Recovers Most HIV Transmission Times and Sources

Finally, we applied *TnT* to the 11-patient HIV transmission chain. In this chain, all but one direct transmission relationship are known from clinical surveys (Lemey et al. 2005; Vrancken et al. 2014). We analyse the genes *gag* and *pol*, with *TnT* allowing us to perform joint analyses by inferring two gene trees and the transmission tree between hosts simultaneously. Table 3 shows the posterior estimates of tree statistics and tree generating parameters. Figure 5 shows the transmission network inferred by *TnT*. We can see that most of the transmission events are inferred correctly and with high probability. All incorrectly determined transmission events can be explained by our knowledge of the data. First, clinical data could not designate whether A or B was the index case in this transmission chain. We also know that patient A transmitted to patient F shortly before being sampled, compared to its own infection date (see Figure S1 in Vrancken et al. 2014). These facts may explain why our model cannot conclusively determine the index case and the direction of A–F transmission. Additionally, *TnT* incorrectly determines the donor for patient H. From clinical data, we know that patient B infected both I and H; and patient I was infected around 2 years before patient H. Patients I and H were sampled only after transmission to both had occurred. A similar bias was observed in determining the transmission of B → I and B → H from single *pol* gene tree analyses by Vrancken et al. 2014.

**Table 3:**
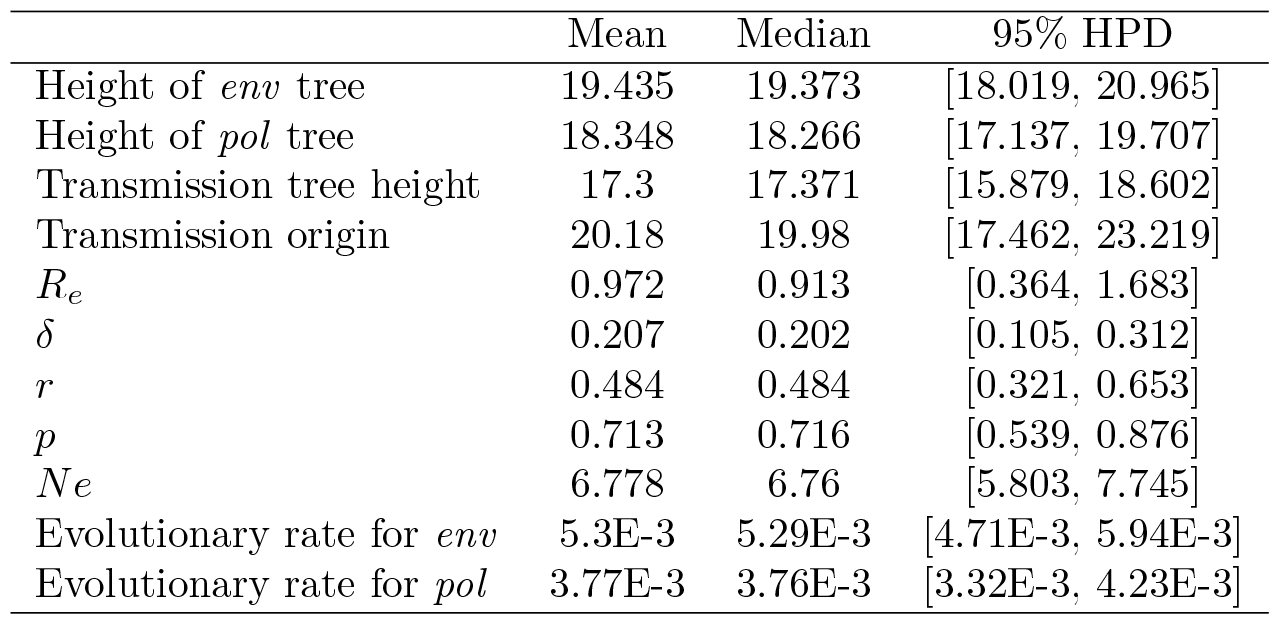
Posterior estimates obtained applying *TnT* to HIV data. The evolutionary rate is measured in substitutions per site per year. HPD – highest posterior density.

|  | Mean | Median | 95% HPD |
| --- | --- | --- | --- |
| Height of *env* tree | 19.435 | 19.373 | [18.019, 20.965] |
| Height of *pol* tree | 18.348 | 18.266 | [17.137, 19.707] |
| Transmission tree height | 17.3 | 17.371 | [15.879, 18.602] |
| Transmission origin | 20.18 | 19.98 | [17.462, 23.219] |
| $R_e$ | 0.972 | 0.913 | [0.364, 1.683] |
| $\delta$ | 0.207 | 0.202 | [0.105, 0.312] |
| $r$ | 0.484 | 0.484 | [0.321, 0.653] |
| $p$ | 0.713 | 0.716 | [0.539, 0.876] |
| $Ne$ | 6.778 | 6.76 | [5.803, 7.745] |
| Evolutionary rate for *env* | 5.3E-3 | 5.29E-3 | [4.71E-3, 5.94E-3] |
| Evolutionary rate for *pol* | 3.77E-3 | 3.76E-3 | [3.32E-3, 4.23E-3] |

**Figure 5.**
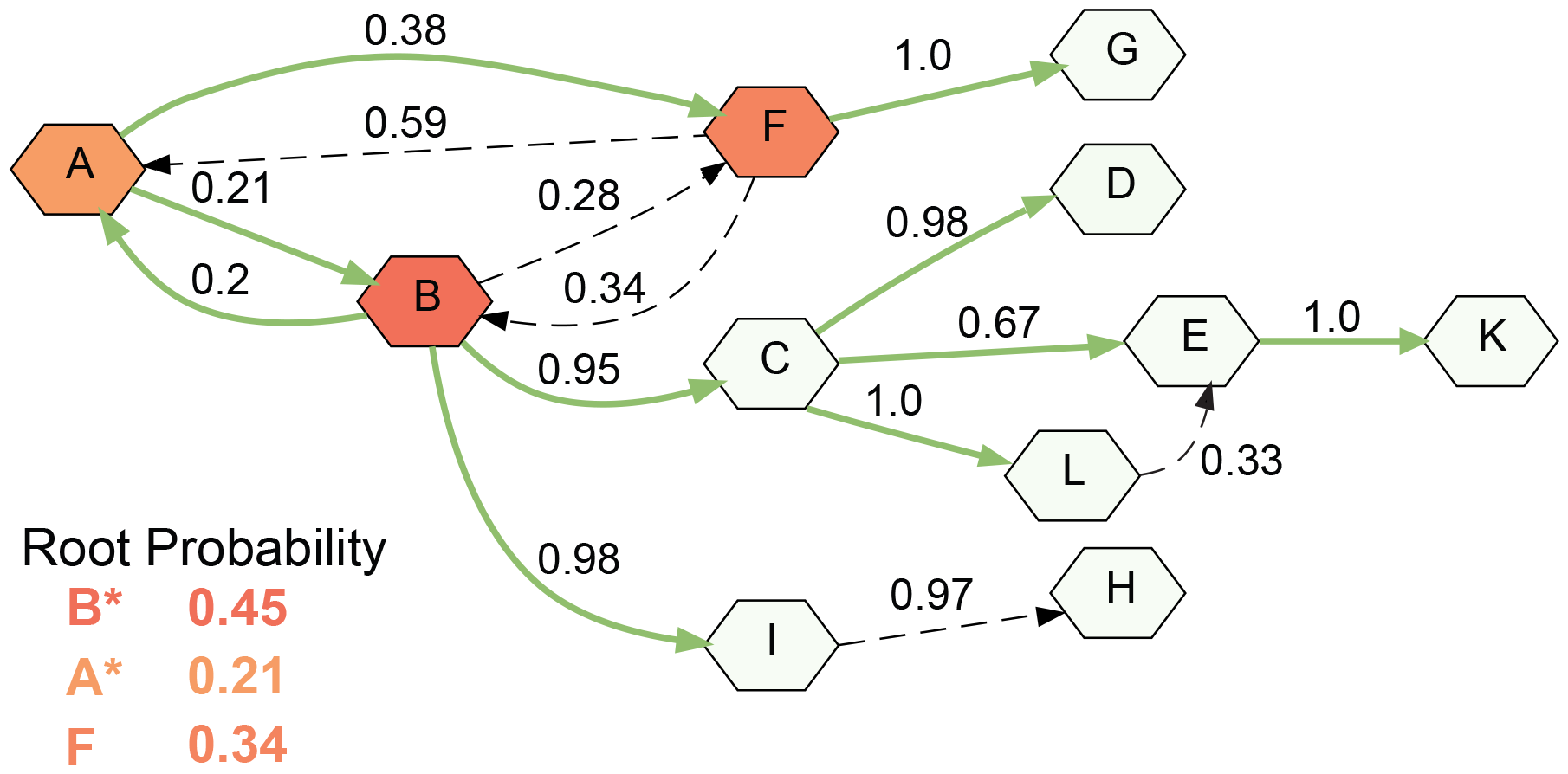
Summary network of the transmission events obtained by *TnT*. Nodes show hosts. Edges show transmission direction and are annotated by posterior probability of directed transmission event. Probabilities below 0.1 are not shown. Edges are green solid lines if they are supported by the clinical information and black dashed lines otherwise. Node color and intensity shows the probability of it being the root. In the table, * denotes possibilities supported by the clinical data.

Figure 3b shows the infection period for each host that was determined in three different ways, namely by clinical surveys, by *TnT* with transmission bottlenecks, and by *TnT* when assuming no transmission bottleneck. Hosts A and B are excluded here due to unknown transmission direction and no estimate for the start of the infection period in clinical data. We see that *TnT* is largely in agreement with clinical data and in some cases (hosts L, K, E, C) the possible infection time window is narrowed. The only very wide time window (host F) is explained by *TnT* not being able to conclusively determine the index case. Notably, the posterior estimates obtained when assuming no bottleneck have much more uncertainty, as evident by wider HPD intervals.

## 4 Discussion

This study introduces the *Transmission Tree (TnT)* model, a method that combines epidemiological and evolutionary processes to generate time-calibrated transmission trees using incompletely sampled data. Utilising a Bayesian approach, *TnT* facilitates a nuanced understanding of transmission dynamics, thus contributing to our knowledge of infectious disease propagation. *TnT* allows for the quantification of transmission times and bottleneck size, even in the cases of incomplete sampling of the outbreak. Additionally, by providing the posterior of congruent trees, *TnT* eliminates the need to reconcile differences or inconsistencies that may arise from individual gene tree analyses. Last, while not shown in this paper, *TnT* is capable of uncorrelated relaxed clock (Alexei J. Drummond et al. 2006) and skyline (Alexei J Drummond et al. 2005; Stadler et al. 2013) analyses, as well as effective population size estimation per transmission tree branch. As we have demonstrated, our new technique provides a robust model-based approach to transmission tree reconstruction and bottleneck strength estimation. That said, there are several ways the method could be improved.

Firstly, while simulations have shown that our stochastic mapping technique is powerful in estimating the number of unobserved transmissions and time of transmission to sampled hosts, the current version does not use all available information from the genetic sequences. While our MCMC implementation samples bottleneck-induced events at unobserved transmissions, our stochastic mapping algorithm ignores these events and only uses the posterior samples of the SABD model rates and transmission tree. An implementation of the stochastic mapping algorithm in which samples are conditioned on events in the gene tree could produce more accurate results.

Second, we assume a single bottleneck at transmission. Bottlenecks have been shown to occur within the host, especially between compartments (Pfeiffer and Kirkegaard 2006; Gutiérrez, Michalakis, and Blanc 2012). For example, there may be additional bottlenecks in the time that transmission occurs via the sexual route, and viral strains have been sampled from blood. As we described in the Methods, the SABD model, currently used as a prior for the transmission tree, does not allow for a set of samples to be assigned to the same host. In case of incomplete sampling, the probability of unobserved transmission events between such samples may be small, but it is not exactly zero. Although our implementation of the rejection prior will keep such samples on the same transmission tree lineage, if genetic sequences support within host bottlenecks, the probability of a bottleneck between same host samples may be very high. Since unobserved transmissions are allowed, we may sample bottleneck-induced events on the gene trees. This may bias the posterior of pairwise coalescence probability at a bottleneck. This issue could be addressed by applying the birth-death stratigraphic ranges model instead of SABD as the transmission tree prior (Stadler et al. 2018). This model, currently more known for macroevolution applications, allows assigning a set of samples to the same taxa and correctly does not marginalise over the unobserved branching events between these samples. This prior, together with tree operators aware of sample-to-taxon assignment, would remove the bias above and potentially speed up the convergence of *TnT*. However, it has not yet been implemented in BEAST2 or equivalent Bayesian phylodynamics software.

Third, *TnT* makes several trade-offs. To allow individual gene trees and ensure their congruence with the transmission tree, complex MCMC operators are necessary. Together with a rejection prior, it makes *TnT* much more computationally costly than *SCOTTI*. Furthermore, an instantaneous bottleneck implies that no mutation occurs during it, which means that this model can only be applicable to very slow evolving pathogens or those where the bottleneck time is assumed to be vanishingly small in comparison to the infection duration. Further work could be focused on allowing for time-dependent effective population size functions.

Ultimately, *TnT* is a detailed approach that explicitly accounts for the epidemiological and evolutionary process on a calendar time scale. By intregrating these complex dynamics, *TnT* is capable of producing time-calibrated transmission trees, even when based on incompletely sampled data.

The intricacy of the *TnT* model architecture, while a strength, also renders it slower compared to simpler models. The computational demand means that—in its current form—*TnT* is best suited for analyses of smaller datasets. Looking forward, we envision *TnT* not only as a direct tool for research, but also as a possible benchmark for developing novel heuristics, approximations, and methodologies aimed at larger datasets. As the amount of pathogen sequencing data available for analysis continues to grow, we show that robust statistical tools can use it to uncover detailed epidemiological information.

## 4.1 Data and Software Availability

The *TnT* package for BEAST2 is open source and freely available at https://github.com/jugne/TnT. The code for all analyses can be found at https://github.com/jugne/TnT-material. HIV data available from GenBank, accession numbers KF773157—KF773625.

## 4.2 Acknowledgement

U.S., T.G.V. and T.S. thank ETH Zürich for funding. U.S., T.G.V. and T.S. received funding from the European Research Council (ERC) under the European Union’s Horizon 2020 research and innovation programme grant agreement No 101001077 (PhyCogy).

