## Supplementary Materials for "Integrating Transmission Dynamics and Pathogen Evolution Through a Bayesian Approach"

April 14, 2024

### 1 Supplementary information

#### 1.1 Gene Tree Prior

Given a birth-death process with birth rate $\lambda$, death rate $\mu$, sampling through time rate $\psi$ and removal probability $r$. Here, all rates are positive and $\lambda \neq \mu$. We know Stadler 2010; Stadler et al. 2012; Gavryushkina et al. 2014 that the probability that an individual alive at time t before today has no sampled extinct or extant descendants is:

$$P_0(t) = \frac{\lambda + \mu + \psi + c_1 \frac{e^{-c_1 t}(1-c_2)-(1+c_2)}{e^{-c_1 t}(1-c_2)+(1+c_2)}}{2\lambda}$$

where

$$c_1 = |\sqrt{(\lambda - \mu - \psi)^2 + 4\lambda\psi}| \text{ and } c_2 = -\frac{\lambda - \mu - \psi}{c_1}.$$

Define

$$q(t) = \frac{4}{2(1 - {c_2}^2) + e^{-c_1 t}(1 - c_2)^2 + e^{c_1 t}(1 + c_2)^2}.$$

We may also extend the model to include incomplete sampling of extant hosts with rate $\rho$. Then,

$$c_2 = -\frac{\lambda - \mu - 2\lambda\rho - \psi}{c_1}.$$

#### Integrating Over Non-Observed Transmission events

Here, a phylogenetic tree generated by birth-death process, with the rates above, produces a oriented transmission tree. Every bifurcation on this tree is a transmission event. The tree is best visualized as asymmetric tree where lineage that started before the bifurcation event and continues afterwards is a *donor* and the lineages that branches from it at the bifurcation event is *recipient*. Following Bokma, Brink, and Stadler 2012, the chance of unobserved bifurcation is $2\lambda P_0$. The expected number of hidden transmission events on a branch between origin time $t_o$ and end time $t_e$, when time increases backwards, is

$$E[S] = 2\lambda \int_{t_e}^{t_o} P_0 dt = (t_o - t_e)(\mu + \psi + \lambda - c_1) + 2ln \left[ \frac{(c_2 - 1)e^{-c_1 t_e} - c_2 - 1}{(c_2 - 1)e^{-c_1 t_o} - c_2 - 1} \right]. \quad (1)$$

The probability that a individual at time point $t_o$ is still infectious, has not been sampled, did not have any observed transmission events by the time $t_o - t_e$ and that the unobserved transmission events happen at time points $T_1, T_2, ..., T_S$ is

$$e^{-(\mu+\psi+\lambda)(t_o-t_e)} \frac{(2\lambda)^S}{S!} P_0(T_1) P_0(T_2) \cdots P_0(T_S).$$

As noted in Bokma, Brink, and Stadler 2012, to obtain the probability of $S$ unobserved transmission events in the interval $(t_o, t_e)$, we integrate over $T_1, T_2, ..., T_S$ and condition on the probability that the branch is observed during this interval $P_1(t_e)/P_1(t_o) = q(t_e)/q(t_o)$ (see Stadler 2010, but note our definition of $q(t)$ are inverse):

$$P(S) = e^{-(\mu+\psi+\lambda)(t_o-t_e)} \frac{P_1(t_e)}{P_1(t_o)} \frac{(2\lambda)^S}{S!} \left( \int_{t_o}^{t_e} P_0(T) dT \right)^S \quad (2)$$

where

$$\begin{aligned} \frac{P_1(t_e)}{P_1(t_o)} &= \frac{2(1 - {c_2}^2) + e^{-c_1 t_o}(1 - c_2)^2 + e^{c_1 t_o}(1 + c_2)^2}{2(1 - {c_2}^2) + e^{-c_1 t_e}(1 - c_2)^2 + e^{c_1 t_e}(1 + c_2)^2} \\ &= \frac{2(1 - {c_2}^2)e^{-c_1 t_o} + e^{-2c_1 t_o}(1 - c_2)^2 + (1 + c_2)^2}{2(1 - {c_2}^2)e^{-c_1 t_e} + e^{-2c_1 t_e}(1 - c_2)^2 + (1 + c_2)^2} \cdot \frac{e^{c_1 t_o}}{e^c_1 t_e} \\ &= \left[ \frac{(1 - c_2)e^{-c_1 t_o} + c_2 + 1}{(1 - c_2)e^{-c_1 t_e} + c_2 + 1} \right]^2 \cdot \frac{e^{c_1 t_o}}{e^c_1 t_e} \\ &= \left[ \frac{(c_2 - 1)e^{-c_1 t_o} - c_2 - 1}{(c_2 - 1)e^{-c_1 t_e} - c_2 - 1} \right]^2 \cdot e^{c_1 (t_o - t_e)} \end{aligned}$$

Then, full expression of equation (2) is

$$\begin{aligned} P(S) =& e^{-(\mu+\psi+\lambda-c_1)(t_o-t_e)} \left[ \frac{(c_2 - 1)e^{-c_1 t_o} - c_2 - 1}{(c_2 - 1)e^{-c_1 t_e} - c_2 - 1} \right]^2 \\ & \times \frac{(t_o - t_e)(\mu + \psi + \lambda - c_1) + 2ln \left[ \frac{(c_2-1)e^{-c_1 t_e} - c_2 - 1}{(c_2-1)e^{-c_1 t_o} - c_2 - 1} \right]}{S!}, \end{aligned}$$

which is equal to

$$P(S = s) = \frac{E[S]^s}{s!} e^{-E[S]}. \quad (3)$$

Further, we want to know the chance of an unobserved transmission event that *change the host of a branch*. That is, until unobserved transmission event, the host associated with this branch is the donor and immediately after this event it is the recipient. We can show that number of such events between time $t_o$ and $t_e$ follows a Poisson distribution with mean $\frac{E[S]}{2}$:

$$
\begin{aligned}
P[X=x] &= \sum_{s=x}^{\infty} P[S=s] \left(\frac{1}{2}\right)^{s} \binom{s}{x} \\
&= \sum_{s=x}^{\infty} \frac{E[S]^{s}}{s!} e^{-E[S]} \left(\frac{1}{2}\right)^{s} \binom{s}{x} \\
&= \sum_{s=x}^{\infty} e^{-E[S]} \left(\frac{E[S]}{2}\right)^{s} \frac{1}{x!(s-x)!} \\
&= e^{-E[S]} \left(\frac{E[S]}{2}\right)^{x} \frac{1}{x!} \sum_{s=0}^{\infty} \left(\frac{E[S]}{2}\right)^{s} \frac{1}{s!} \\
&= e^{-E[S]} \left(\frac{E[S]}{2}\right)^{x} \frac{1}{x!} e^{\frac{E[S]}{2}} \\
&= e^{-\frac{E[S]}{2}} \left(\frac{E[S]}{2}\right)^{x} \frac{1}{x!}
\end{aligned}
$$

.

### Bottleneck strength definition

We define the instantaneous bottleneck with strength expressed as the amount of coalescent time $\tau$ it would take for the coalescence to happen at a usual rate. Coalescent time is scaled by effective population number N ($t$ generations is $T = \frac{t}{N}$ coalescent time units). Alternatively, we may define bottleneck strength as a probability for two lineages to coalesce: $p := 1 - e^{-\tau}$.

Note, that assumption of an instantaneous bottleneck is only valid if the period that is being modelled is significantly longer than the actual duration of the bottleneck.

### Probability for $i$ lineages derive from $j$ lineages in the past

Here we describe some previous results that will be useful when deriving the gene tree likelihood. From equation (6.1) in Tavaré 1984, probability that current $i$ lineages descend from $j$ lineages at $T$ coalescent time in the past is

$$
g_{ij}(T) = \begin{cases} \sum_{k=j}^{i} \frac{(2k-1)(-1)^{k-j} j_{(k-1)} i_{[k]}}{j!(k-j)! i_{(k)}} e^{-\frac{k(k-1)}{2}T}, & \text{if } 1 \leqslant j \leqslant i \\ 0, & \text{otherwise} \end{cases} \tag{4}
$$

where $a_{(k)} = a(a+1)\dots(a+k-1)$ for $k \geqslant 1$ with $a_{(0)} = 1$ and $a_{[k]} = a(a-1)\dots(a-k+1)$ for $k \geqslant 1$ with $a_{[0]} = 1$. To obtain a valid probability distribution, require also $\sum_{j=1}^{i} g_{ij}(T) = 1$, for all $T$.

Note, that probability $p$ from the previous section is simply a special case: $p = g_{21}(\tau) = 1 - e^{-\tau}$.

### Some combinatoric factors

Number of ways for $i$ to coalesce into $j$ Rosenberg 2003:

$$
I_{ij} = \binom{i}{2}\binom{i-1}{2}\dots\binom{j+1}{2} = \frac{i!(i-1)!}{2^{i-j} j!(j-1)!} \tag{5}
$$

Define a coalescent cluster as $i+1$ lineages coalescing into 1. Let us have two simultaneously evolving coalescent clusters. We are interested in all possible orderings of the coalescent events within and between these clusters. Note that lineage from one cluster cannot coalesce with a lineage from another cluster. Number of ways $i+1$ lineages coalesce into 1 and $j+1$ lineages coalesce into 1 simultaneously Rosenberg 2003:

$$W_{ij} = \binom{i+j}{i} \tag{6}$$

We can extend this to any number $z \in \mathbb{N}$ of coalescent clusters. Define the set $\mathbf{S} = \{s_k | s_k \in \mathbb{N} \wedge s_k > 0 \wedge 1 \leq k \leq z\}$, where $s_k + 1$ is the number of lineages in each cluster. Then

$$W_{s_{1:z}} = \binom{s_1 + s_2 + s_3 + ... + s_z}{s_1} \binom{s_2 + s_3 + ... + s_z}{s_2} \dots \binom{s_{z-1} + s_z}{s_{z-1}} \tag{7}$$

### Gene tree likelihood

Let $g$ be the gene tree from the gene trees space $G_i$ over the $i^{th}$ alignment. Going backwards in time along the transmission tree branch, we can note all events on this embedded gene tree and their respective times $t_0, t_1, \dots, t_n, t_{n+1}$. Times $t_0$ and $t_{n+1}$ are, respectively the start and end of the branch b. Equivalently to Heled and Drummond 2009 we denote the gene tree lineage history over branch $b$ $L_b(g) = \{l_m, t_m | 0 \leq m \leq n+1\}$. Here $l_m$ is the number or lineages at the time $t_m$. In contrast with Heled and Drummond 2009, at the number of lineages at the time $t_{n+1}$ depends not only on bifurcation and sampling events, but also on possible higher order coalescence, introduced by the hidden transmission events and transmission bottleneck. Also, if the branch $b$ corresponds to a donor host on a transmission tree, there are no coalescent events (neither bifurcating, nor multifurcating) on a gene tree $g$ at the time $t_{n+1}$. Otherwise, if the branch $b$ corresponds to a recipient host on a transmission tree, we model an instantaneous transmission bottleneck at time $t_{n+1}$. Note, that bottleneck increases a probability for two coalescent lineages, going backwards in time. It however, does not imply that any order of coalescence will be necessarily observed.

We define gene tree likelihood as product of the contributions for bifurcating and higher order coalescent events, allowing for multiple mergers, as well as intervals between these events and observed transmission contribution. Excluding end times $t_0$ and $t_{n+1}$, there are two possible events that contribute to the likelihood: coalescent event and hidden transmission event. Following gene tree topology, each bifurcation can account for a coalescent event or a coalescent event that is a result of transmission bottleneck at a hidden transmission event. Each higher order coalescence can only be a result of instantaneous bottleneck and therefore accounts for a hidden transmission event. It is important, that hidden transmission event may also be a part of an interval between the bifurcating and multifurcating coalescent events. This is the case when there are no are no coalescence, induced by a transmission bottleneck at this hidden transmission event.

As derived above, the chance that there is a hidden transmission event at time $t_m$ is $\lambda P_0(t_m)$. Denote $N_b$ the effective population size over the transmission tree branch $b$, inverse of which is the coalescent rate. Finally, we need the probability of $i$ lineages reducing to $j$ in bottleneck duration coalescent time $\tau$, $g_{ij}(\tau)$, number of ways $i$ lineages can coalesce to $j$, $I_{ij}$, and number of ways to order coalescent events in case of multiple mergers, $W_{s_{1:z}}$. Instantaneous transmission bottlenecks may result in multiple mergers, which, in turn, can consist of multifurcations. We can look at each separate multifurcation as coalescent cluster, defined above. There are many ways to order coalescent events within and between these clusters to obtain the same instantaneous multiple merger. For $z$ multiple mergers, each consisting out of $s_k + 1$, $k \in \{1, 2, ..., k\}$, lineages which coalesce into 1, we can order such coalescent events in $W_{s_{1:z}}$ ways. Within each cluster we have $I_{s_k+1,1}$ ways to coalesce.

Then, the total number of ways to obtain a multiple merger with $z$ coalescent clusters is:

$$W_{s_{1:z}} \prod_{i=1}^{z} I_{s_k+1,1} \tag{8}$$

Probability of one specific coalescent path of $i$ lineages coalescing into $j$ lineages in coalescent time $\tau$ we have:

$$\frac{1}{I_{ij}} g_{ij}(\tau) \tag{9}$$

Excluding start and end of the transmission tree branch $b$, multiple mergers and multifurcations on a gene tree can only be the result of non-observed transmission events. Then contribution of such event, where $i$ lineages are reduced to $j$ lineages, and every $k^{th}$ merger reduces $s_k + 1$ lineages to 1, is:

$$W_{s_{1:z}} \prod_{k=1}^{z} I_{s_k+1,1} \frac{1}{I_{ij}} g_{ij}(\tau) \lambda P_0(t) \tag{10}$$

Bifurcating coalescent on a gene tree might be a result of a normal coalescent process at rate $\frac{1}{N_b}$ or a unobserved transmission bottleneck. Then, contribution of bifurcating coalescent event is:

$$\frac{1}{N_b} + \frac{1}{I_{i,i-1}} g_{l,l-1}(\tau) \lambda P_0(t) \tag{11}$$

Interval contribution between events is:

$$e^{-\int_{t_0}^{t_1} \frac{\binom{l}{2}}{N_b(t)} dt} e^{-\int_{t_0}^{t_1} (1 - g_{l,l}(\tau)) \lambda P_0(t) dt}, \tag{12}$$

where first exponential term stands for no coalescent events happening in this interval and the second term stands for no unobserved transmission events that result in any order coalescence on the tree.

We now define a helper function, which notes the probability of the observed coalescence pattern at a bottleneck:

$$h(\tau, i, j, z, \mathbf{S}) = W_{s_{1:z}} \prod_{k=1}^{z} I_{s_k+1,1} \frac{1}{I_{ij}} g_{ij}(\tau).$$

Gathering the above equations, we may write the probability of gene tree lineage history over a transmission tree branch $b$, by factoring it into the interval contributions $V_m$ and event contributions $E_m$:

$$P(L_b(g)|N_b, \tau, \lambda, \mu, \psi) = \prod_{m=1}^{n+1} V_m \cdot E_m.$$

Here,

$$V_m = e^{-\int_{t_{m-1}}^{t_m} \frac{\binom{l_m}{2}}{N_b} dt} e^{-\int_{t_{m-1}}^{t_m} (1 - g_{l_m, l_m}(\tau)) \lambda P_0(t) dt}$$

and

$$E_m = \begin{cases} \frac{1}{N_b} + h(\tau, l_m, l_m - 1, 1, 1) \lambda P_0(t_m), & \text{if event } m \text{ is a bifurcation, } m < n+1; \\ h(\tau, l_m, l_{m+1}, z, \mathbf{S}) \lambda P_0(t_m), & \text{if event } m \text{ is a higher order coalescence, } m < n+1; \\ h(\tau, l_m, l_{m+1}, z, \mathbf{S}), & \text{if } b \text{ is a recipient branch and } m = n+1; \\ 1, & \text{if } b \text{ is a donor branch and } m = n+1. \end{cases}$$

#### 1.2 MCMC moves

Here we briefly describe new and adjusted tree operators.

##### Gene Tree Operators

Gene tree operators were severely influenced by *pitchfork* BEAST2 package (Vaughan 2023). Notably, any gene tree is embedded within a transmission tree and restricted by it. Any move that breaks this embedding will not be proposed or will be rejected (depending which one is easier and computationally fatser to implement).

**CreateMergersOrReheight** This operator selects a non-leaf node $i$ which may be a coalescent node or a polytomy root. If it is not the tree root, it may also be part of a multiple merger event. We define the interval between the time of this node's closest child and parent as $[h_c, h_p]$. The operator takes as input a new multiple merger probability $p_m$ and proposes one of the following actions:

- with probability $p_m$ selects another gene tree node $j$, with height $h_j \in [t_c, t_p]$, and changes the height of original node so that $h_i = h_j$. This creates or expands a multiple merger;
- with probability $(1 - p_m)$ chooses a new height the node $j$ from the uniform distribution $Uniform(h_c, h_p)$.

**ExpandCollapseOperator** This operator selects a node $i$ and with equal probability proposes one of the actions:

- Let $p_i$ be a parent node of $i$ and $j$ a sister a bi-or multi-furcating sister node of $i$ (i.e. degree of $j \geq 3$) with heights $h_j > h_i$. Collapse the edge $\{i, p_i\}$ and make $j$ the new parent of $i$ thus increasing degree of $j$ by 1.
- Let $p_i$ be a multifurcating parent node of $i$ (i.e. degree of $p_i \geq 4$) and $Gp_i$ a parent node of $p_i$. Draw a new height from uniform distribution $Uniform(h_{p_i}, h_{Gp_i})$ and create a node at this height with parent $Gp_i$ and two child nodes $i$ and $p_i$.

**SPROperator** This is a subtree prune and regraft operator for a gene tree with bottleneck induced events. It takes a probability $p_b$ of attaching the root edge of the pruned subtree at an existing height of remaining tree. That is it may attach at a bi- or multifurcating node, increasing the degree of this node by one, or creating a new node of degree 3 at the same height of another node or nodes in the tree, thus creating a multiple merger event. With probability $1 - p_b$, the operator behaves as the usual subtree prune and regraft.

**SubTreeSlideOperator** This is a subtree slide for a gene tree with bottleneck induced events. It takes two probabilities, $p_1$ and $p_2$, with $p_1 + p_2 \leq 1$. With probability $p_1$, the new attachment after the slide move is at a existing node, creating or expanding a polytomy. With probability $p_2$, the attachment after the slide move is at the same height as another non-leaf node, creating or expanding a multiple merger. With probability $1 - p_1 - p_2$, the usual subtree slide move is performed.

**TransmissionAttach** This operator selects a transmission event on the transmission tree and performs one of the two actions:

- Select a non-root gene tree node $i$ from the gene tree nodes which are embedded in the transmission subtree, rooted at the recipient side of this transmission event and which has a parent node $p_i$ with height greater than the transmission event height. Parent node $p_i$ may not be a polytomy or part of a multiple merger. Prune the subtree rooted at $i$ and re-attach at the transmission event height (leaving the same sibling and parent nodes for $p_i$).
- Select a non-root gene tree node $i$, with parent at the same height as the transmission event. Let $Gp_i$ be the parent of $p_i$. Reheight $p_i$ drawing the new height from the uniform distribution $Uniform(h_{p_i}, h_{Gp_i})$.

#### Transmission Tree Operators

**SwapOrientations** This operator takes two child nodes of a coalescent (transmission) event on the oriented tree, where left child is a donor and right child is the recipient, and swap their orientations (as well as donor-recipient roles represented by these branches).

We have also made necessary changes to the *CoordinatedExponential* and *CoordinatedUniform* operators from the StarBeast2 package (Ogilvie, Bouckaert, and Drummond 2017), narrow *ExchangeOperator* and *ScaleOperator* from BEAST2 core (Bouckaert et al. 2019) and *LeafToSampledAncestorJump*, *SAUniform*, *SAExchange* and *SAWilsonBalding* from sampled-ancestors package (Gavryushkina et al. 2014), in order to properly address polytomies and multiple mergers resulting from the instantaneous events on a gene tree.

### 2 Supplementary Tables and Figures

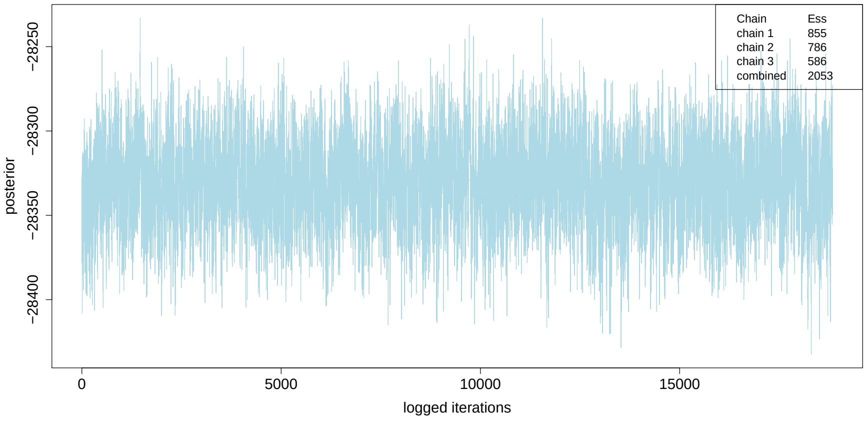


(a)

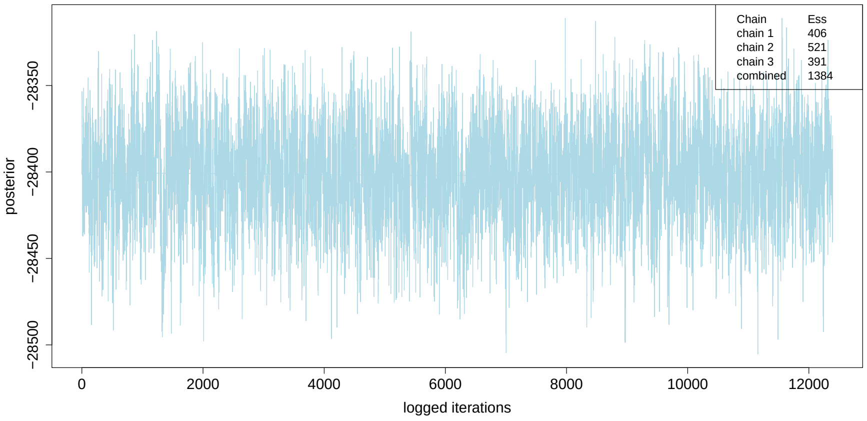


(b)

Figure 1: **Posterior trace for combined chains obtained by *TnT*.** (a) Estimating the bottleneck, 3 chains, combined after 30% burnin. (b) Bottleneck strength set to 0, 3 chains, combined after 20% burnin. ESS–effective sample size.
